# Deciphering drug response and phenotypic heterogeneity of cancer cells using gene ensembles of regulatory units defined by chromatin domains

**DOI:** 10.1101/2023.01.15.524115

**Authors:** Neetesh Pandey, Madhu Sharma, Arpit Mathur, Chukwuemeka George Anene-Nzel, Muhammad Hakimullah, Priyanka Patel, Indra Prakash Jha, Omkar Chandra, Shreya Mishra, Jui Bhattacharya, Ankur Sharma, Roger Foo, Kuljeet Sandhu, Amit Mandoli, Ramanuj DasGupta, Vibhor Kumar

## Abstract

The effect of co-localization of genes in the topologically associated domains (TADs) and their activity as a regulatory unit in cancer samples and cells, together with drug-response, needs comprehensive analysis. Here, we analyzed the activity of TADs using cancer-cell transcriptomes along with chromatin-interaction and epigenome profiles to understand their relationship with drug-response. Our analysis of 819 cancer cell-line transcriptomes revealed that their response to multiple drugs was more correlated with the activity of individual TADs than genes. Applying our approach to 9014 cancer patients’ data (20 different cancer types) also revealed a higher association between survival and the activity of thousands of individual TADs in comparison to their genes. CRISPR-mediated knock-out of regulatory sites inside a TAD associated with cisplatin-response of oral cancer cells and discovery of primate-specific gain of synteny of genes within a TAD containing EGFR gene and its contribution towards cancer malignancy demonstrate greater utility of TAD-activity based analysis.

## Introduction

The orchestration of chromatin organisation and the activity of different genomic segments are important factors for the transition of states and the response of cells. Several studies have been dedicated to linking properties of cancer cells and chromatin organization ^1^, structural variation in their genomes ^2^. Even though genetic mutation and escape from apoptosis are involved in the development of a cancerous state, regulatory changes in epigenome and gene expression always become instrumental in such transition^3^. Cancer therapy tends to spare some cancer cells, even when they lack resistance, causing mutations ^4^, and several studies have tried to find reasons for such a phenomenon. A scientific group involved with Cancer Cell Line Encyclopedia (CCLE) attempted to estimate response for drugs ^5^ using mutation and expression profiles. Many attempts have been made to provide non-genetic explanations like reprogramming and stemness, which are linked with increased xenobiotic resistance and the efflux pump expression^6,7^, greater DNA repair effectiveness^6^, resilience and stress response ^8,9^. The heterogeneity of tumors with the capacity of a few cancer cells to transition states has also been explained as a cause for drug-resistance^10^. In every scenario, chromatin organization plays a crucial role in the growth of cancer cells and their drug-resistant versions ^11,12^.

Chromatin is organized in the form of multiple domains, often called topologically associated domains (TAD). TADs can also be labeled as naturally defined genomic regions segregated from other genomic loci ^13,14^. TADs have been proposed to act as functional regulatory units^15,16^ and reported to be largely stable across cell types ^13,17^. In fact, a few studies have shown the conservation of approximately 54% of TAD boundaries in homologous regions in the genomes of humans and mice ^13^. The gene co-localization also exhibits a pattern driven by cellular and evolutionary mechanisms, such as a gene-dense locus 3p21.31 on chromosome 3, which is known to harbor the largest cluster of tumor suppressor genes ^18^. The co-localisation and synteny of genes involved in similar or related functions could be due to multiple reasons, such as avoiding the recombination of beneficial alleles in a locus^18^ or the co-expression. Salem *et al.* have shown that paralogs are more often in the same TAD than random pairs of genes ^19^. The mutation rates in TADs and their boundaries have been studied in the context of the development of cancer ^20^. However, multiple factors could be involved in the formation and strengthening of TAD boundaries, such as CTCF, G4 quadruplexes ^21^, and repeats ^22^. Barutcu *et al.* have shown that deleting CTCF-rich regions near TAD boundaries does not necessarily disrupt the TADs ^23^. Similarly, Akdemir et al.^20^, reported that only 14 percent of TAD boundary deletions resulted in change of expression of proximal genes by two-fold. Hence, it is not trivial to trace the effect of a mutation on TAD-boundary maintenance or disruption. Therefore, the abnormality in the activity of genes inside a TAD could be a more direct indication of the effect of the TAD boundary. Moreover, there are many questions on TADs and their effect on the regulation of cancer cell properties and their drug-response, as well as the utility of TADs for translational medicine (see Supplementary Notes). To resolve such problems and to find answers to several questions, we developed TADac, an approach to estimate TAD-activity as an enrichment of expression of a set of genes in a TAD to find their link with drug response and cancer malignancy.

First, we analyzed the chromatin interaction landscape for head and neck squamous cell carcinoma (HNSCC) cell lines to comprehend the pattern of activity of TADs in drug resistance. Later, we expanded our study by analyzing the transcriptome of 819 cell lines reported in the CCLE database and the response for 544 drugs reported in the Cancer Therapeutic Response Portal (CTRP)^24^. Further, we used our approach to draw inferences from transcriptome profiles of 9014 patients across 20 cancer types made available by the Cancer Genome Atlas (TCGA)^25^ consortium.

## Results

The integrative analysis using our unique approach to chromatin interaction profiles and single-cell transcriptomes of patient-derived HNSCC (or HNSCC) cell lines laid the foundation for analyzing larger transcriptome data-sets of cell lines from the CCLE database and tumors from TCGA databases.

### Oral cancer Hi-C profiling, TAD patterns, and their association with drug resistance

We profiled the chromatin-interaction landscape of HNSCC cell lines using the Hi-C protocol. Specifically, we used patient-derived primary cultures (PDCs) of HNSCC established by Chia *et al.* ^26^, which include primary HNSCC cells of two patients’ primary tumors (HN148P, HN137P) and metastatic cancer cells of patient HN120 (HN120M) (Supplementary Figure S1). We also performed ChIP-seq using antibodies for H3K27ac and H3K4me3 histone modifications in HN137P and their metastatic version HN137M cell lines. For the single-cell transcriptome, TADac calculates TAD gene-set enrichment for every single-cell ^27^ (see Methods). To calculate TAD-activity using the bulk expression profile, it uses the approach of Gene-set variation analysis (GSVA)^28^. A brief outline of the work is also shown in Figure 1A.

**Figure 1:**
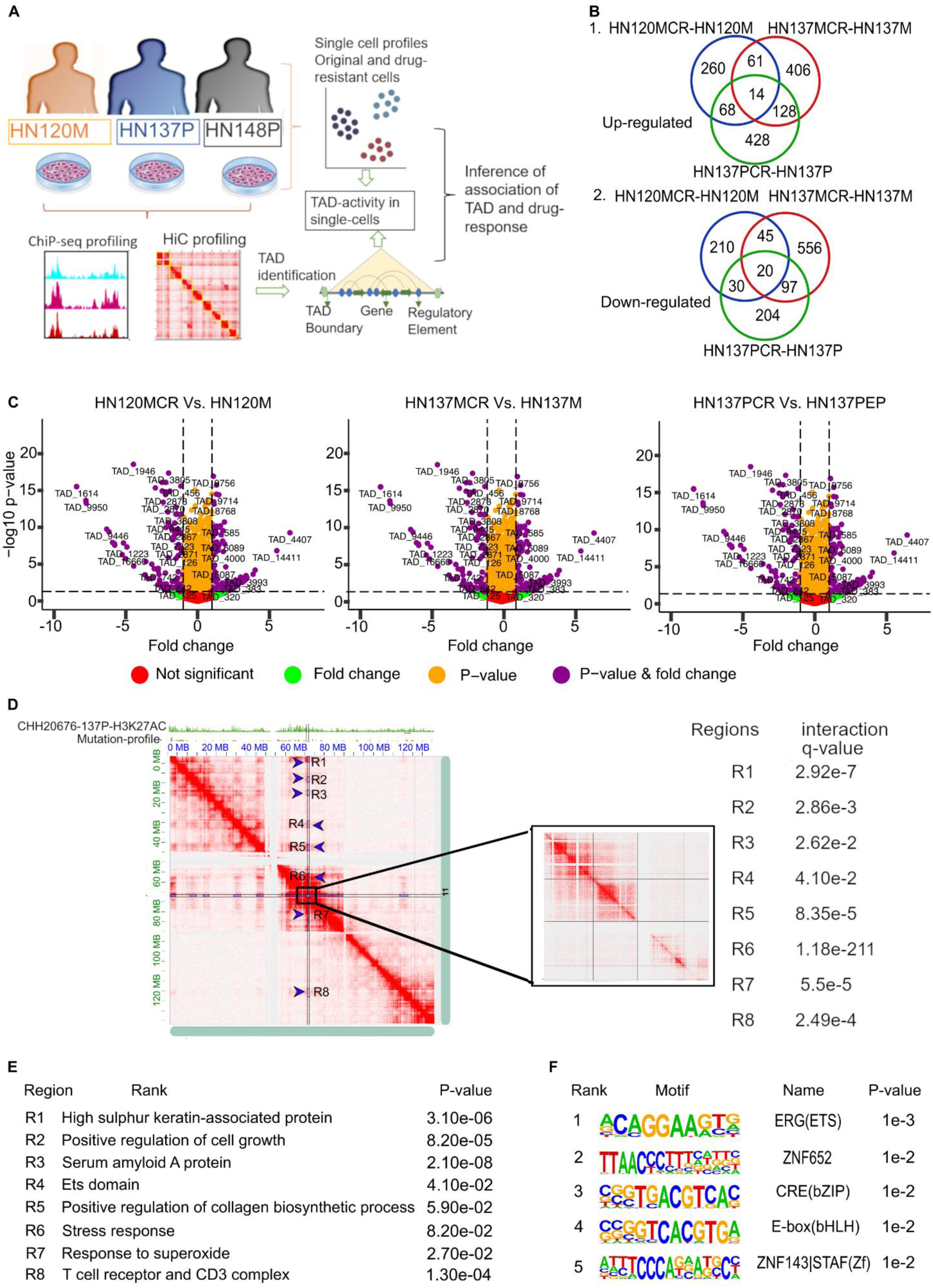
Analysis of chromatin interaction profiles and domain activity of head and neck cancer patient-derived cell lines. **A)** A description of the profiling and analysis of the chromatin interaction profiles of three head and neck squamous cell carcinoma (HNSCC) cell lines: metastatic cell line HN120M from patient HN120, primary tumor cells HN137P and HN148P from patients HN137 and HN148, respectively. The union list of TAD boundaries derived from Hi-C profiles of HNSCC cell lines was used for downstream analysis. **B)** Venn diagrams showing the number of TADs commonly up-regulated or down-regulated in cisplatin-resistant versions of HNSCC cell lines. **C)** The fold-change and significance (P-value) of the TAD-activity between HNSCC cell lines and their cisplatin-resistant versions are depicted on the volcano plots. **D)** The chromatin interaction visualization for chromosome 11 using the Hi-C profile of 137P cells. The TADs (in the 11q13.3-4 bands) downregulated in all three HNSCC cell lines is the present region inside the rectangle indicated in the middle. Other regions with distal interaction with the TAD in 11q13.3-11q13.4 bands are shown with arrows as R1, R2, R3 … R8). The q-values (or adjusted P-values) for interaction among 11q13.3-4 TAD region and R1.. R8 are also shown. **E)** The enriched gene-ontology terms of genes present in different regions interacting with 11q13.3-11q13.4 bands. **F)** The enriched motifs in H3K27ac peaks in down-regulated TAD in 11q13.3-11q13.4 band and R1, R2 …. R8 region.

#### TAD-activity alterations with cisplatin resistance development in HNSCC cells

We made a union list of TADs detected using the Hi-C profile of HNSCC cells and calculated differences in their TAD-activity level among oral cancer cell lines and their cisplatin-resistant variants. For this purpose, we used the single-cell expression profiles of HNSCC cell lines HN137P (paired-end), HN137M, HN120M, and their cisplatin-resistant versions HN137PCR, HN137MCR and HN120MCR published by Sharma *et al.*^10^ and further confirmed for correctness using mutational profiles ^29^. Overall, 14 TADs had consistent upregulation (P-value < 0.05, fold change > 1.5) in activity in cisplatin-resistant versions of all three HNSCC cell lines (HN137PCR, HN120MCR, HN137MCR) (Supplementary Table 1). Among 14 TADs, three TADs are from the 13q13.1 band and harbor genes associated with actin filament reorganization, mitotic sister chromatid cohesion, and nucleotide excision repairs like BRCA2, FRY, and PDS5B. Other commonly upregulated TADs in other regions also include genes associated with mitotic sister chromatid cohesion, response to DNA damage, TGF-beta activation, drug transport, and mitochondrial translational elongation known as signatures of resistance and response to cisplatin (see Supplemental File-1).

A total of 20 TADs had consistently down-regulated activity in cisplatin-resistant versions of multiple cell lines (HN137PCR, HN120MCR, HN137MCR) (Figure 1B-C, Supplementary Table 2). Since we allowed overlap among TAD regions to accommodate for TAD-hierarchy, eight out of 20 commonly downregulated TADs lay within region chr11:70150000-71470000 (hg19) in 11q13.3 and 11q13.4 bands and showed some hierarchy. For brevity, we call them 11q13.3-4 TADs. All the 11q13.3-4 TADs have 5 known keratin associated protein genes (KRTAP5-7, KRTAP5-8, KRTAP5-9, KRTAP5-10, and KRTAP5-11) along with DHCR7 and NADSYN1 genes. Out of eight 11q13.3-4 TADs, the longer TADs also contain SHANK2, CTTN, and a few more genes with both known and unknown functions. The copy number amplification and over-expression of genes in amplicon in 11q13-11q14 bands have been reported to be present in malignant tumors of many patients with HNSCC and other cancer types ^30^, however its regulation in response towards cisplatin has rarely been reported.

Visualization of the chromatin interaction landscape of HNSCC cells in chr11 revealed that the genomic region in 11q13.3-11q13.4 bands has statistically significant connections (q-value < 0.05) to a few other regions in the same chromosome (Figure 1D, regions R1, R2 R3, R4, R5, R6, R7, R8) (supplemental Methods). Such intensity of interaction of 11q13.3-4 bands and R1-R8 regions was not found in other cell types, like cardiomyocytes and astrocytes (Supplementary Figure S2). The 11q13.3-4 TADs could be part of a larger regulatory condensate in HNSCC cells. In addition, to 11q13.3-4 TADs, (with genes KRTAP5-7, KRTAP5-8, KRTAP5-9, KRTAP5-10, KRTAP5-11) and the region R1 connected to them also has many genes of family KRATAP5 (KRTAP5-1, KRTAP5-2, KRTAP5-3, KRTAP5-4, and KRTAP5-5). The regions R2, R3, and R4 have significant enrichment (P-value < 0.05) for the gene-set of ‘positive regulation of cell growth’, ‘central carbon metabolism in cancer’, and ETS family genes, respectively, which are known to promote cancer cell proliferation. The region R7 has enrichment for folate receptor activity, which is known to be upregulated in oral cancer (Figure 1E) ^31^. The H3K27ac ChIP-seq peaks (in HN137 cell lines) present in 11q13.3-4 TADs and regions connected to them (R1-R8) had enrichment of motifs for transcription factors (TFs) EWS: ERG, FLI1, and ETV2, which belong to the ETS family (Figure 1F) and are known to be involved in promoting cancer ^32,33^. Translocation could also be involved in causing higher interaction of 11q13.3-4 TADs with regions R1, R2, and R8 compared to the background. Nevertheless, our approach of analysis of HNSCC cells’ profiles revealed important TADs in 11q13.3-11q13.4 bands lying at the hub of chromatin interaction of regions which could be involved in HNSCC cell proliferation and could be important targets for influencing the viability of HNSCC cells.

### Generalization of the relationship between drug sensitivity and TAD-activity in cell lines from multiple cancer

The insights gained through analysis using HNSCC cell lines encouraged comprehensive investigation of drug-response in a wider range of cancer cell lines using the same approach. Hence, we compiled a union list of TADs detected in our HNSCC cell lines, and TADs reported in the TADKB database ^14^. Our approach is based on reports by multiple researchers on the conservation of TAD boundaries across cell types ^13,34^, which we further verified in cancer cells (Supplementary Figure S3A-B). Thus we used a common union list of TADs to calculate TAD-activity scores for 819 CCLE cell lines^5^ and correlate their TAD-activity with the pIC50 value of drugs reported for respective cell lines at CTRP^24^ (see Methods). Compared to natural TADs, the null-model TADs with random boundaries had a lower correlation between their activity level and drug pIC50 values (Supplementary Figure S4A-B). For HNSCC cell lines, there were high correlation values between the activity of pIC50 values of many drugs and the activities of many TADs, which we clustered into 8 groups (see Figure 2A). The clustering results revealed clusters of TADs whose activity had both a positive correlation with pIC50 values for a group of drugs and a negative correlation with pIC50 values for another group of drugs (Figure 2A). However, cluster-1 and cluster-5 (Figure 2A) had a negative correlation with the pIC50 value of most drugs and were enriched for genes involved in the positive regulation of peptide serine and antigen processing by MHC, respectively. We repeated the same analysis using transcriptome profiles of CCLE cell lines of LUSC (Lung squamous cell carcinoma) and BRCA (Breast cancer)(Supplementary Figure S5). Clustering of TADs based on correlation with pIC50 values of drugs reveals not only major functional categories in the context of drug-response but also has the potential to guide better drug combinations.

**Figure 2:**
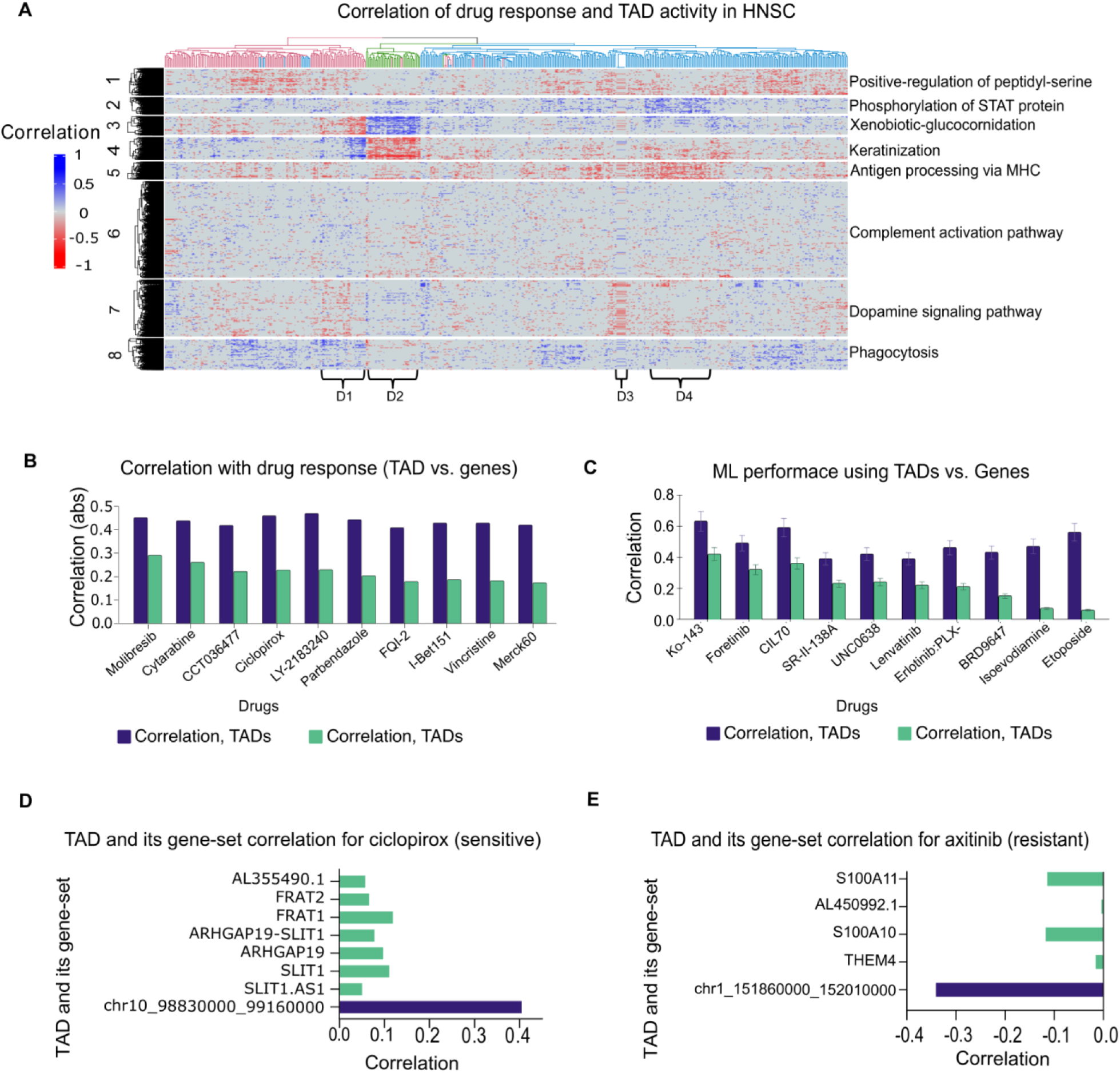
Pattern of correlation between TAD-activity and drug response. **A)** Heatmap-based illustration for correlation among TAD-activity and pIC50 value for drugs in head and neck squamous cell carcinoma (HNSCC) cell lines from CCLE database. Here row resepent TADs and column represent drugs. Only those rows (TADs) and columns (drugs) are present where absolute value of correlation is above 0.30. Four sub-clusters of drugs are also highlighted here. As shown, cluster-3 TAD-activity had a negative correlation with pIC50 values of drugs in the D1 group but positive correlations with drugs in the D2 group. Drugs of group D4 had a pIC50 value negatively correlated with the activity of TADs in cluster-5 and positively correlated with cluster-2 TADs. **C)** Barplot of highest absolute correlation values achieved for a few drugs between their pIC50 level and individual TAD activity (blue) or single gene expression (green), using cell lines of all cancer-types in CCLE database. **C)** The correlation between actual and predicted pIC50 values using a machine learning model (using CCLE database transcriptomes) are shown in a bar chart. Here the features were either TAD-activity or gene expression. For many drugs, even when numerous genes are used, the prediction of pIC50 values (the effectiveness or accuracy) was noticeably lower than the value attained by TAD-activity score. **D)** Correlation with PIC value for drug (Ciclopirox) with activity of a TAD (blue) (chr10_98830000_99160000) across and its respective genes. As shown, none of the genes individually have as high a correlation with pIC50 value of Ciclopirox as the activity of their host TAD. **E)** Comparison of correlation values of activity of a TAD (blue) (chr1_151880000_152010000) and expression of genes in its respective gene-set with pIC50 value of axitinib across available cell-lines of all cancer-types.

Using cell lines for all cancer types in the CCLE database together, resulted in lower correlations values between TAD-activities and pIC50 of drugs (Supplementary Figure S6). Thus it indicates that the association of TADs with the response towards drugs has specificity for cancer cell type. Nevertheless, there were many TADs whose activity had a notable correlation with drug response even when cell lines of all types of cancers in the CCLE set were used ^35^. Clustering correlations between TAD-activity and pIC50 values for drugs for multiple cancer-types in CCLE set and ontology enrichment revealed keratinocyte differentiation, extracellular matrix organization, cornification, aging and glucuronidation as common functions enriched for TAD associated with multi-drug resistance (Supplemental File 2, Supplementary Figure S6). Whereas ‘negative regulation of IL-6’ appeared as an enriched term for genes in TADs with sensitivity towards multiple drugs. However, there were a few drugs which had a high correlation between their pIC50 values and the activity of only one or two TADs, highlighting specificity. Such as drug 1,9-Pyrazolonthrone had a negative correlation (below −0.30, P-value < 0.01) only with activity of TAD at chr20:43940000-44410000 and pIC50 value for drug Erastin had significant correlation (below −0.3, P-value < 0.01) with TAD at chr17:18590000-18710000 (Supplemental File 3).

While using all cancer cell lines of the CCLE database for associating TAD-activity with drug response, we found an interesting pattern: for many drugs, the correlation between the activity of a TAD and the pIC50 values was substantially higher than the best correlation for any single-gene (Figure 2B). Such results highlight the fact that even if genes involved in response to a drug could vary in different types of cancer, the mechanism would remain the same. For example, many paralogous genes with similar functions often co-localise in TAD. We found that for many drugs, even when we used many genes in a machine learning model to predict pIC50 values, the accuracy achieved is much lower than using prediction based on the activity of natural TAD gene sets (Figure 2C). In fact, for many TADs, none of their own genes had a comparable correlation between expression and pIC50 value (Figure 2D-E) with the same drug (Supplementary Figure S7A-B). Such as for the drugs Ciclopirox, LY-2183240 Lenvatinib, Axitinib, and Foretinib, the association between their pIC50 values were substantially greater with TAD-activity than any of their genes (Figure 2D-E, Supplementary Figure S7). Such results indicate that the enrichment score of activity of a gene-set of TADs could be more effective in estimating response than using single genes for many drugs.

### TAD-activity patterns and their influence on malignancy were revealed through patient profiles in TCGA PAN-cancer

There could be multiple reasons for the variation of activity of genes in a TAD in tumors, such as i) change in the regulation of gene activity, ii) mutation at the TAD boundary, or genomic structural variation^36^. iii) Copy number variation (CNV) of the genomic region consisting of a TAD. Nevertheless, their substantial effect on the activity of genes could be tracked using the TAD-activity score. Hence, we studied variations in TAD-activity using the cancer sample transcriptomes provided by the TCGA consortium, using the same union list of TAD as used for the analysis of the CCLE data-set. The differential activity of TADs among tumor versus normal samples and their clustering revealed multiple TADs with cancer-specific activation (upregulation). A few groups of TADs such cluster-12 and cluster-11 in Figure-3A, also showed upregulation in multiple types of cancer. The clusters of TADs made using their differential activity could not highlight any particular biological process with significant enrichment (FDR < 0.1), except for cluster-11. Keratinocyte differentiation and cellular glucuronidation terms were enriched by genes in cluster-11. Such results indicate that genes in a large number of TADs with perturbed activity in cancer of different types do not associate with specific biological processes.

Further, we focused on TADs associated with survival in 20 different types of cancer (see Methods and Supplementary Figure S8). The number of TADs associated with survival in 2 and 3 cancer types was substantially more than TADs with survival association in only one cancer type (Figure 3B). To confirm top survival-associated TADs without the effect of conditional dependencies, we used Bayesian network modeling where every TAD was represented by a node (see Methods) (Supplementary Figure S9). The TAD in the 11q13.3-4 region was among the top 30 TADs with a direct influence on survival in HNSCC (TCGA-HNSC) patients according to Bayesian network modeling of survival (Supplementary Figure S9A). Kaplan-Meier plots for a few TADs in the context of CESC, LUSC, and HNSCC cancer are provided in supplementary figures S10 and S11. Analysis of hazard ratios revealed that many TADs could have significant associations (P-value < 0.05) with survival in multiple cancers, and their effects could vary based on the context of cancer-type (Figure 3C).

**Figure 3:**
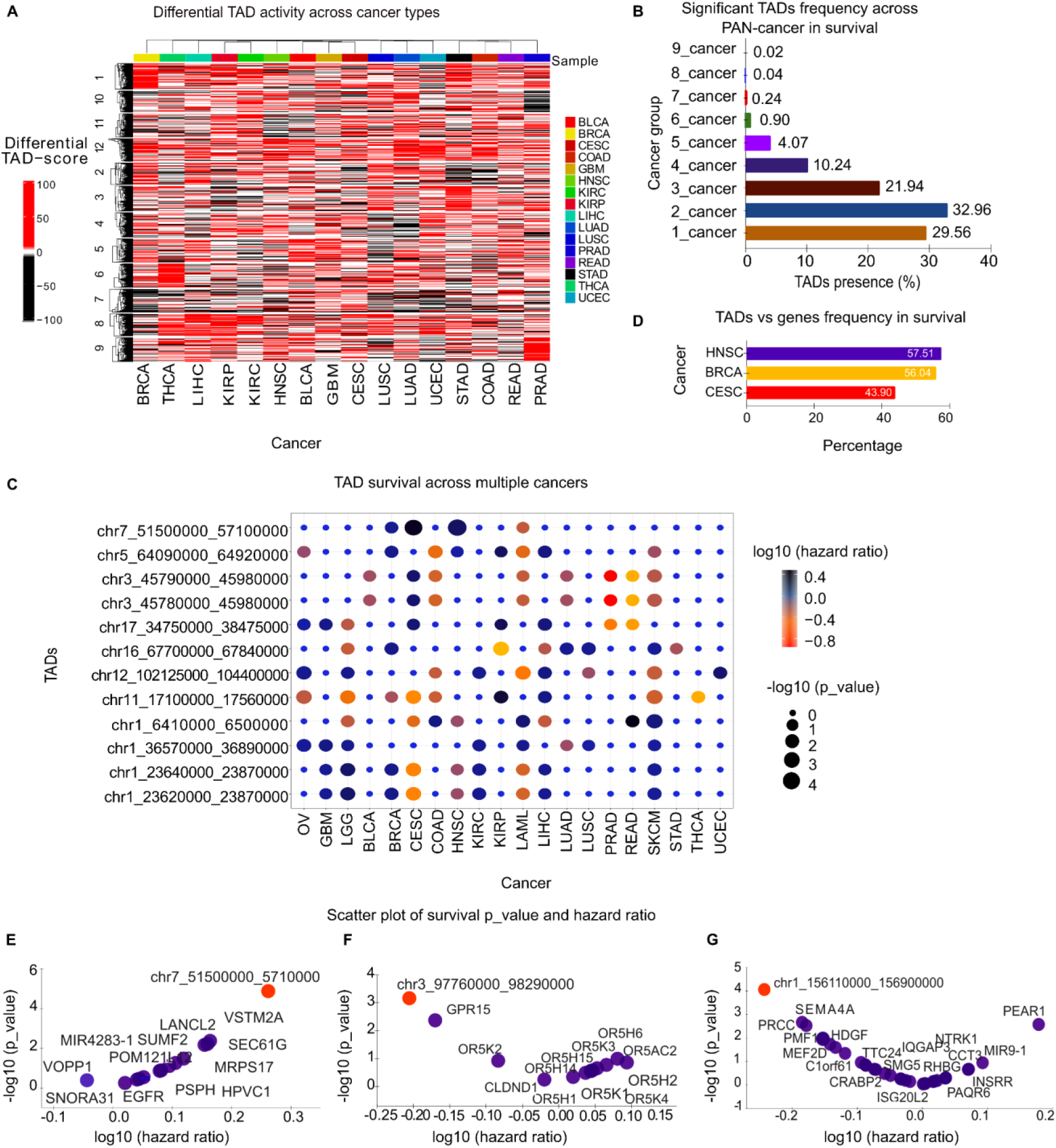
Activity of TADs across multiple cancer TCGA samples and their association with survival. **A)** Heatmap of fold changes of median TAD-activity (tumor Vs normal sample) across 16 cancer types, calculated using TCGA data. **B)** Here, the proportion of numbers of TADs associated with survival in only one or 2 or 3 cancer types are shown. For association, the P-value threshold was 0.05, and proportion was calculated by division by the number of TADs that had an association with survival in at least one cancer type. **C)** dot-plot showing P-values and hazard ratios for survival association with TADs across PAN-cancer. **D)** Percentage of TADs having more influence on association with survival than any of their genes. The percentage was calculated with respect to all TADs that had an association with survival in the given cancer type. **E)** TAD (chr7_51500000_57100000) and its genes survival association P-value in TCGA-HNSC patients are illustrated using scatter plots. **F)** TAD (chr3_97760000_98290000)-gene survival P-value comparisons (in TCGA-HNSC) are illustrated using scatter plots. **G)** TAD (chr1_156110000_156900000)-gene survival P-value comparisons (in TCGA-HNSC) are illustrated using scatter plots.

There could be a possibility that a TAD’s association with survival could be an artifact of the effect of a few genes, which could be independently more influential for survival than the activity of its host TAD. However, we found that for multiple TADs, their activity had a higher influence on survival than any of the genes inside them. The fraction of such TADs with more influence on survival than any of their genes were 57%, 56%, and 44% in HNSC, BRCA, and CESC cancer, respectively (Figure 3D). Such as a TAD containing EGFR gene located at chr7:51500000-57100000 has a more significant association (P-value < 1E-4) between its activity and survival than any of its genes in HNSC (Figure 3E, Supplementary Figures S10, S11), BRCA, and CESC. Other such examples for TCGA-HNSC are TADs located at chr3:97760000-98290000 and chr1:156110000-156900000, which have more association with survival than any of the genes lying within their boundaries (Figure 3F-G, Supplementary Figures S10). Overall, TAD-activity-based survival analysis hints about the effect of co-localisation of genes and their coordination for major biological processes in causing cancer malignancy.

We evaluated coherence of association of survival outcomes in cancer patients with TAD-activity and their correlation with drug response in cancer cell lines (see Supplementary Methods). We sorted the TADs associated with survival (P-value < 0.05) in TCGA-HNSCC patients according to the number of drugs with which they had absolute correlation values higher than 0.3 for HNSCC cell lines in the CCLE database. The top 50 TADs (survival-associated, P-value < 0.05) providing resistance (negative correlation with pIC50; r <-0.3) towards the highest number of drugs had a significantly higher hazard ratio (wilcoxon Rank Sum, P-value < 0.01) than the top 50 TADs correlated with sensitivity towards the maximum number of drugs (see Figure 4A). The same pattern was observed when top 10 or top 100 TADs were used. Out of the top 10 TADs (with survival P-value < 0.05) correlated towards resistance with the highest number of drugs (r < −0.3 with pIC50), 50% had hazard ratios above 1.25. Whereas none of the top 10 TADs (with survival P-value < 0.05) correlated for sensitivity towards the highest number of drugs (r > 0.3 with pIC50) had a hazard ratio above 1.25 (Figure 4B). A similar trend was observed for the top-50 or top-100 TAD categories (see Figure 4C). Such results highlighted coherence between influence of TAD on drug-response in cell lines and malignancy in patients as well as useful TAD based markers associated with both cancer severity and multi-drug resistance.

**Figure 4:**
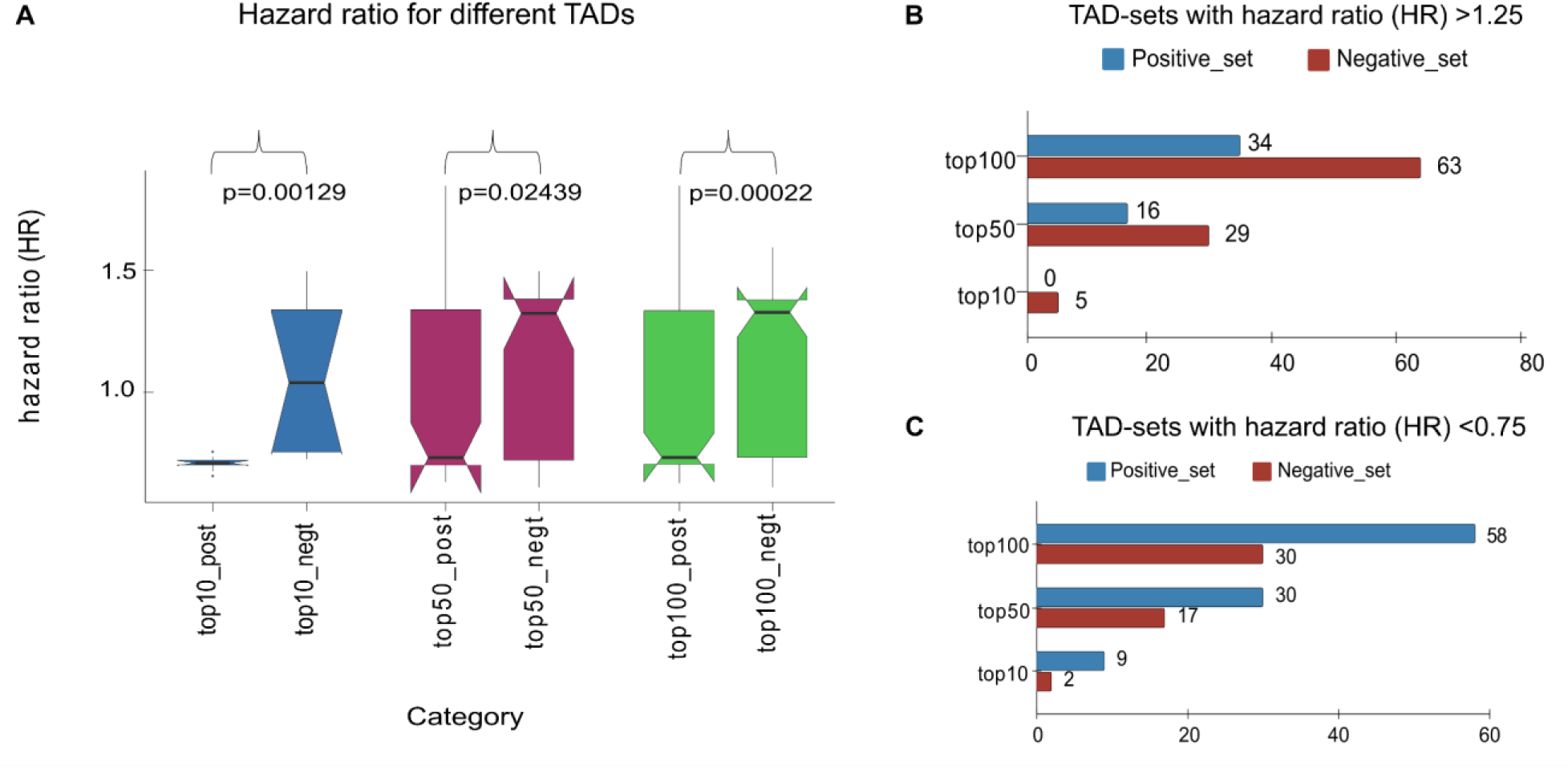
Coherence between the association of TAD-activity with survival among TCGA patients and relationship between TAD and drug-response in CCLE cell lines. **A)** hazard-ratio distribution of TADs correlated with drug-response, across the top 10, 50, and 100 categories. Here top10_post contains those TADs, which are 10 most frequently positively correlated (R > 0.3) with pIC50 values of drugs. Similarly, top10_neg contains those TADs which have the top 10 highest frequency of negative correlation with drug pIC50 value. Top50_post, top50_negt, top100_post, and top100_negt groups are on the same trend. The hazard ratio for the activity of TADs negatively correlated with pIC50 value (or positive correlation with IC50) was above one (low survival). Similarly, the inverse is true for positively associated TADs. **B)** Frequency for TADs in different groups (top50_pos or top50_neg) for hazard ratio cutoffs greater than 1.25 (in TCGA cohort). **C)** Frequency of TADs in different groups, for hazard ratio less than 0.75.

### Case study of TADs in 11q13.3-11q13.4 region

We focused on 11q13.3-4 TADs as they appeared to be at the center of a hub of chromatin interaction in chromosome 11 in HNSCC cells and located on the genomic site with high copy number amplification in the HNSCC tumor of multiple patients in TCGA cohort (Figure 5A). Hi-C profile of HNSCC cells around other TADs with top association with survival, did not show strong distal interactions similar to 11q13.3-4 region (Supplementary Figure S12). The association between the enrichment of the KRTAP5 gene-set and TCGA-HNSC (HNSCC) patient survival was insignificant (P-val=0.054). Moreover, the promoters of KRTAP5 family genes in 11q13.3-4 TADs did not have H3K27ac or H3K4me3 histone modifications at their promoters in HN120M, HN137P and HN137M HNSCC cell lines (Supplementary Figure S13A). The CNV of the gene-set in 11q13.3-4 TADs is significantly (P-value < 0.05) associated with the survival of HNSCC patients (Figure 5C). The synteny of genes in 11q13.3-4 TADs appear to be mostly preserved across most of the mammals (Figure 5D) (see Methods) and point towards a possible conservative force for the co-localization of genes (Supplementary Figure S14A). Two locations with high chromatin-accessibility in tumor samples of TCGA-HNSC (or HNSCC) patients overlap with eRNA in HNSCC samples, as reported by the TCGA consortium (Figure 5E) (Supplementary Figure S13B). CRISPR-based knockout using gRNA for those two locations decreased the viability of HNSCC cell line Cal27. However, CRISPR-based knockout using the same guide RNA in control cells (HEK293) did not have any effect (Figure 5F). RT-qPCR results showed increased expression of FADD, DHCR7, CTTN, RNF121, and PPFIA1 genes in comparison to housekeeping gene (GAPDH) in Cal27 cells, which indicated that CRISPR-KO of selected regulatory sites induced cell death. In control HEK293 cells, the genes FADD, DHCR7, RNF121, CTTN, and PPFIA1 did not show any change in expression upon CRISPR-KO of selected regulatory sites (Figure 5G). Such results indicate that identifying important TADs can lead to finding relevant enhancers to be targeted to affect the viability of cancer cells.

**Figure 5:**
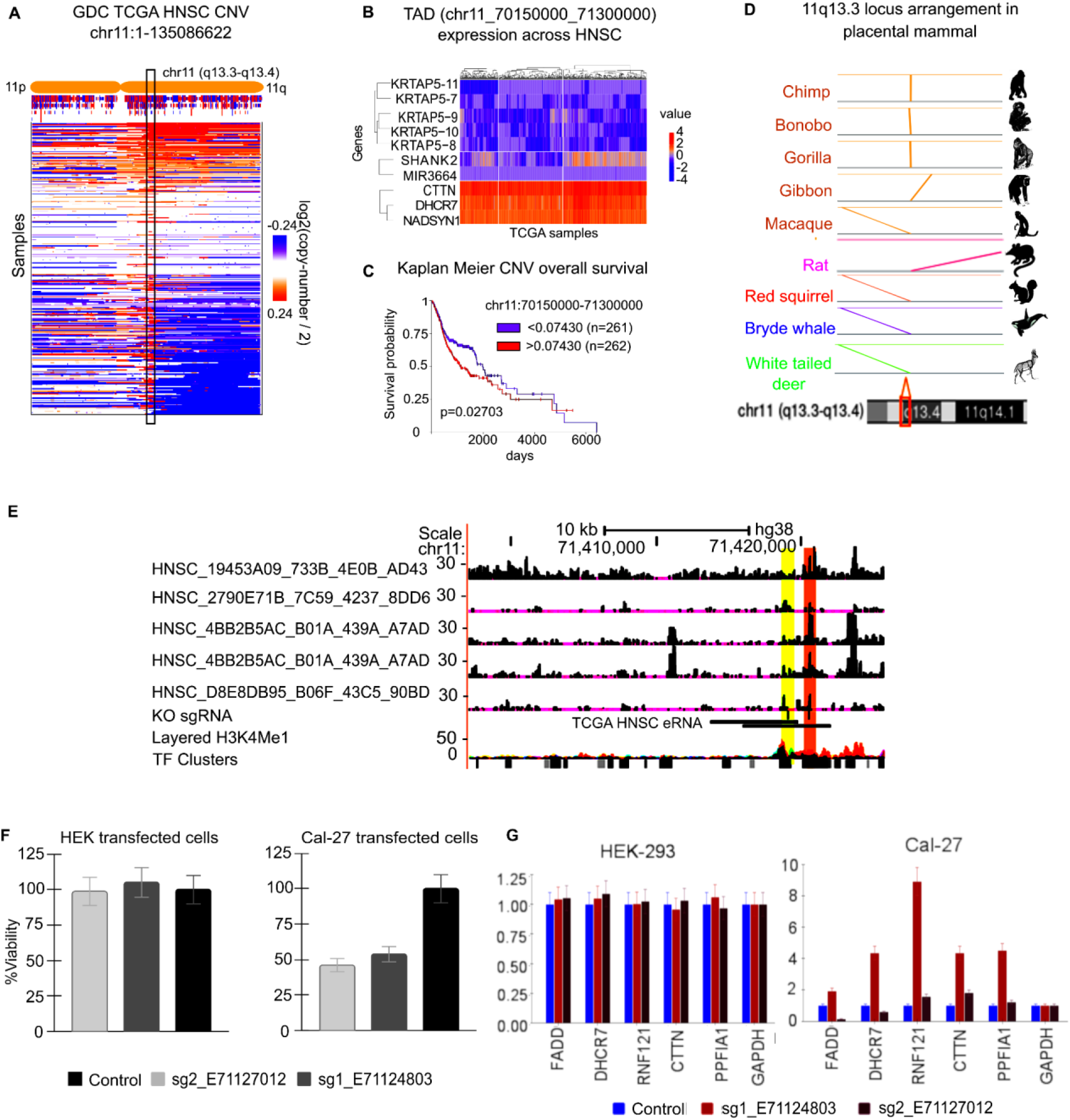
Case study of 11q13.3-4 TAD for identifying and validating target relevant regulatory sites in cancer with CRISPR. **A)** A view of copy number variations of genomic region in chromosome 11 in Tumor samples of Head and Neck Cancer (HNSCC) patients from the cancer genome atlas (TCGA-HNSC). The box shows the regions in 11q13.3 - 11q13.4 bands, which also have high number of CNV in multiple HNSCC patients. **B)** Heatmap of expression of genes in TAD (hg19:chr11-70150000_71300000) in 11q13.3-11q13.4 bands in TCGA-HNSC (HNSCC) samples. **C)** Kaplan Meier (KM) plot showing association of copy number variation (CNV) in genomic location chr11-70150000_71300000, made using Xena browser **D)** A view of conserved synteny of genes in TAD (hg19:chr11-70150000_71300000) of 11q13.4-11q13.4 band across 9 mammalian species. For synteny across 69 mammals, see supplementary Figure S14A. **E)** The UCSC-browser based view of open-chromatin profile (ATAC-seq) of 5 tumor samples of HNSCC patients from TCGA consortium. **F)** Bar plots showing the viability of two cell lines (Cal-27, HEK293) after CRISPR-KO of non-coding sites (enhancers) overlapping TCGA-HNSC-eRNA in a TAD in 11q13.3-11q13.4 band. **G)** RT-PCR based quantification of expression of genes (FADD, DHCR7,CTTN, RNF121, PPFIA1, GAPDH) after CRISPR based knock out of two sites overlapping TCGA-eRNA in TAD (hg19:chr11-70150000_71300000) of 11q13.3-11q13.4 band. The names of gRNAs shown here (sg1_E71124803, sg2_E71127012) also contain their respective locations (hg19, chr11:71124803, and chr1171127012) corresponding to their 5-prime ends. The CRISPR-KO based results are shown for HNSCC cell line (Cal-27) and a control cell line (HEK293).

### Case study of understanding evolutionary and functional pressure in TAD with EGFR gene

The higher influence of activity of TAD at chr7:51500000-57100000 on survival than its gene EGFR in multiple cancer types, raised questions regarding the importance of co-localisation and the function of other genes it hosts. For brevity, we call the TAD located at chr7:51500000-57100000 as EGFR-TAD. The EGFR-TAD tends to have a high copy number in tumors of a few TCGA-HNSC (HNSCC) patients (Figure 6A). We found that genes in EGFR-TAD gained synteny in apes and old-world monkeys even if they remained non-syntenic throughout other mammals and new-world monkeys (Figure 6B, Supplementary Figure S14B). The gain of synteny of genes in apes and old world monkeys in EGFR-TAD hints about their collaborative effect in a common cellular mechanism involved in the development of the brain and severity of cancer, which we further investigated. The effects of CNVs of member genes of EGFR-TAD on survival matched with EGFR gene in multiple types of cancer (Figure 6C), including HNSCC and glioblastoma. It hints about the high probability of co-amplification and co-activity of genes in EGFR-TAD, which was also observed in HNSCC cell lines histone modification pattern (see Supplementary Figure S15). We found that EGFR-TAD has a high enrichment of genomic variations associated with serine and glycine measurements in blood, brain volume and glioblastoma (Figure 6D). A gene PSPH in EGFR-TAD is involved in endogenous serine production through aerobic glycolysis (Supplementary Figure S16) needed by cancer cells^37^. The high enrichment of GWAS (Genome-wide association study) mutations associated with brain volume and gain of synteny in primates with larger brains implies primate-specific adaptive selection on the locus, though at the cost of increased cancer susceptibility.

**Figure 6:**
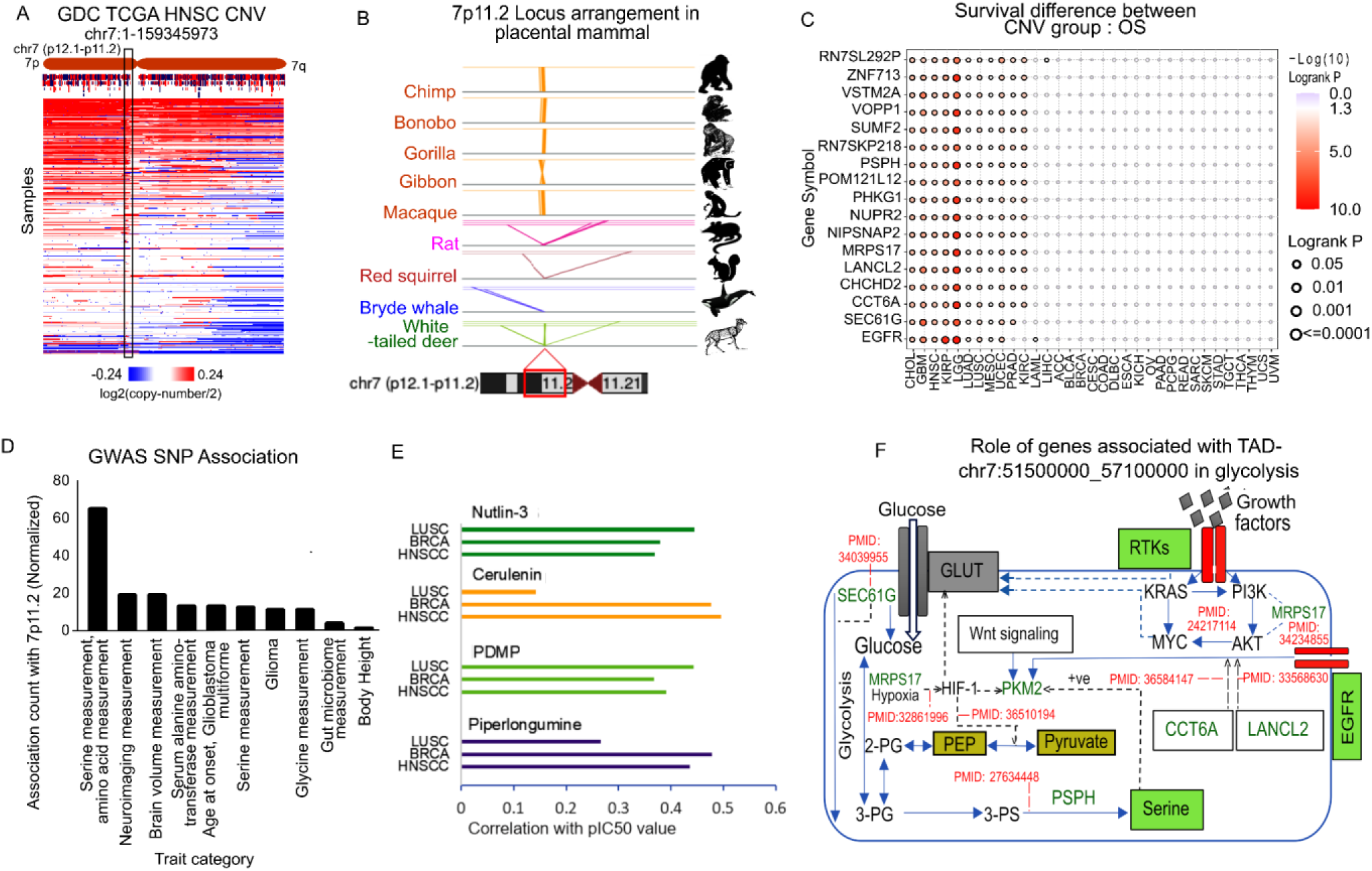
Detail case study for topologically associated domain (TAD) containing EGFR gene. **A)** The visualization of copy number variation (CNV) in chromosome 7 in HNSCC tumor samples in TCGA cohort. The TAD region chr7:51500000-57100000) containing EGFR gene (EGFR-TAD) is shown by a box. **B)** The brief view of conservation of synteny of genes in EGFR-TAD. Here, the state of synteny is shown only for nine mammals, taking humans as reference. For visualizing synteny in 69 mammal species, see (Supplementary Figure S14B) **C)** The association of survival among TCGA-HNSC (HNSCC) patients with CNV of genes in the EGFR-TAD. **D)** Top enriched traits based on presence of GWAS (Genome wide association study) single nucleotide variants present EGFR-TAD. **E)** The correlation of activity of EGFR-TAD with pIC50 value for few drugs in CCLE cell lines are shown. The constraints of numbers of CCLE cell lines allowed such correlation calculation for 3 cancer-type only, namely LUSC, BRCA and HNSCC. The results for drugs with correlation more than 0.35 in at least 2 cancer-types are shown here. **F)** An overview of the involvement of genes in EGFR-TAD in glycolysis, AKT pathway activity and associated processes. The pubmed ID of studies supporting the associations are also mentioned in red color.

Analysis based on CCLE cell-line transcriptomes revealed that drugs with the highest positive correlations between their pIC50 value and activity of the EGFR-TAD affect the AKT pathway or aerobic glycolysis-related processes. In breast cancer cell lines (BRCA), the top 3 drugs with a positive correlation of pIC50 value with EGFR-TAD are piperlongumine, cerulenin, and AZD6482, which have been shown to affect glycolysis directly or indirectly via PI3K/Akt pathway (Supplementary Figure S17 and Supplementary Table 3). Similar trends were observed for HNSCC and LUSC cell lines (Supplementary Figure S17 and Supplementary Table 3). A drug Netulin-3 (MDM2 -inhibitor) (Figure 6E) proven to inhibit endogenous serine production and glycolysis appeared to have sensitivity correlated with the activity of EGFR-TAD (Supplementary Table 3) in all three cancer types (LUSC, BRCA, HNSCC). Multiple genes in the EGFR-TAD support glycolysis and related processes (Figure 6F, Supplementary Table 4), as well as AKT pathway activation. PI3K-AKT pathway is also known to promote aerobic glycolysis^38^. Such results indicate that genes in EGFR-TAD gained synteny in apes to promote cell proliferation in two modes. The first mode is PI3K-AKT pathway activation. The second mode is by supporting aerobic glycolysis for the synthesis of biomolecules (serine, glycine, etc) needed for cellular proliferation. Knowledge of such systematic cooperation of genes can increase the reliability of drug selection based on activity of EGFR-TAD.

## Discussion

Here, we used our computational framework, TADac, to study cancer heterogeneity and drug-response. TAD activity could also be partially reflective of the enrichment of activity of other sets of regulatory elements in a TAD, such as enhancers. The reported low effect of TAD-boundary mutation ^36^ and possibility of many unknown components involved in TAD boundary formation ^23,21^, provide strong reasons in favor of the approach of TADac to use TAD-activity as a direct way to estimate the effect of disruption in TAD-based regulatory units. Moreover, Mohanty et al. ^39^ have shown that copy number and expression variation of genes are not necessarily coupled; hence, the CNV of a TAD region might not be necessarily reflective of its activity. Such findings again highlight the importance of using gene expression directly to estimate the activity of TAD for studying their underlying regulations.

Our analysis clearly indicates that TAD activity could be used as a biomarker for better prediction of response for a few drugs or survival estimation to cope. In many cases, TAD activity could be a better biomarker than any gene for drug-response and survival analysis. The knowledge of dependencies between TAD-activity and drug-response can guide the development of a framework for finding effective drug combinations to prevent redundancies.

The analysis of HNSCC cell lines transcriptomes highlighted TADs in the 11q13.3–4 band, which could cause cisplatin-sensitivity but resistance towards multiple drugs like azacitidine and importazole in oral cancer. The spatial co-localisation of clusters of genes of the KRTAP5 family due to the interaction of 11q13.3 and R1 region could support some major regulatory processes. However, KRTAP5 genes have not been commonly reported for association with properties of cancer cells except by a few anecdotal studies ^40^. Nevertheless, the case study of 11q13.3-4 TADs in squamous cell carcinoma highlighted an alternative approach to reduce search space for the identification of target cis-regulatory elements for cancer therapeutics. Due to the occurrence of copy number amplification of the 11q13.3 band with malignancy ^30^ in many cancer types, there could be wider utility of CRISPR-based knock-out of the identified regulatory region in 11q13.3-4 TAD.

Our result of the discovery of the link between the severity of cancer and an evolutionary force associated with the development of a larger brain in apes also provides a hint about the species-specific effect of associated oncotherapeutics drugs. One such effect could be impact on developing brains in apes. In addition to cell proliferation and drug-resistance, PI3K-AKT pathway and aerobic glycolysis supported by EGFR-TAD has also been associated with immune evasion of tumor cells by modulation of T-cell activity ^41^. Thus, the example of EGFR-TAD also opens a new avenue for analysis of transcriptome using the information of TAD gene-set and search for functional reasons for preservation or gain of synteny of genes to understand the mechanism behind various diseases.

## Methods

### Computational analysis

#### Analysis of Hi-C profile

In our analysis, the HiCUP tool was used to process the Hi-C data. The same pipeline was also used to remove artifacts for downstream analysis ^42,43^.

##### Analysis using the HiCUP pipeline

The HiCUP pipeline involves numerous programs like hicup_truncater, hicup_mapper, hicup_filter, hicup_deduplicator, and hicup_digester which prepares the genome digest file used by hicup_filter. The paired FASTQ files were used to generate Hi-C di-tag paired reads, which were then mapped to the hg19 reference genome. This HiCUP script controlled the pipeline carrying output from one section of the program to the next and executed each script sequentially. The configuration file for the HiCUP program also contains the parameters for the complete pipeline^42^. The majority of reads produced by the HiCUP Mapper script are undoubtedly authentic Hi-C products; a substantial minority are likely not and should be eliminated. For the purpose of finding legitimate Hi-C pairs, the HiCUP filter script is used for the analysis of paired reads along with the file produced by the HiCUP Digester. This might not necessarily be the case, though, as the sequenced region may contain the Hi-C ligation junction. As a result, potentially useful data will be lost when such reads are eliminated from the Hi-C pipeline during the mapping process. By locating ligation junctions among reads and removing sequences downstream of the restriction enzyme recognition site, the HiCUP truncate script helps to fix this ^42^.

##### Pipeline implementation

We used the DpnII digest profile of the hg19 reference genome which was created using hicup_digester for computational analysis. This file represented all possible DpnII fragments in the genome and was used to identify Hi-C artifacts. The alignment process was performed using HiCUP v0.5.10 with Bowtie2 (hg19), and the filtering step’s minimum and maximum di-tag ranges were set to 150 and 1000, respectively. The default settings for the other parameters were used. The final BAM output was produced with a satisfying unique alignment score of more than 65%.

##### Generation of Hi-C contact matrix

We first converted the binary alignment mapped file into arrowhead input format using the samtools view command to create the Hi-C contact matrix ^44^. The normalized Hi-C contact matrix (.hic file) was created using the Juicer tool’s precommand ^45^ with the help of the arrowhead input file. The dump command of Juicer Tools was used to export the observed contact count for each region of the Hi-C map with balanced Knight-Ruiz (KR) normalization. With this, we obtained the text Hi-C matrix (.txt) at 25 kb resolution for all the chromosomes.

#### Detection of topologically associated domains (TADs)

We used the DomainCaller approach from Dixon *et al.* to identify topologically associated domains at 25 kb on all chromosomes. We started with a square contact matrix of the 25 kb Hi-C contact map, and subsequently, directionality indices were created using DI_from_matrix.pl with a bin size of 25,000 and a window size of 125,000. This vector of directionality indices was provided as input for the HMM_calls.m script. Further, we obtained TAD calls by processing HMM via file_ends_cleaner.pl, hmm-state_caller.pl, converter_7col.pl, hmm_probablity_correcter.pl, and, hmm-state_domains.pl. The HMM probability correction script was run with the following parameters as input: min = 2, prob = 0.99, and bin size = 25,000^46^. Secondly, we also curated TAD boundaries from the TADKB database, which gives TADs at resolutions of 50 kb and 10 kb from a variety of cell types, namely human HMEC, NHEK, GM12878, IMR90, K562, HUVEC, and KBM7 ^14,17^. TAD locations for each cell type were obtained with the help of the Directionality Index (DI) ^14^.

#### Analysis of ChIP-seq profiles

Analysis of ChIP-seq profiles was done using DFilter ^47^ to call peaks and make custom tracks. The parameters used for calling peaks were according to the instructions provided on the DFilter help page (https://reggenlab.github.io/DFilter/).

#### TAD gene-sets and calculation of TAD-activity score

TAD gene-sets were made by intersecting the promoter of genes with the TAD boundary location in the hg19 genome version. The activity score of TAD is calculated as enrichment of its gene-set. In the TADac approach, the gene-enrichment score is estimated using the UniPath^27^ approach. The expression of genes in the form of FPKM (fragments per kilobase per million reads) or RPKM of every single cell is converted to p-value, assuming the log-normal distribution. The p-values of non-zero expression of genes belonging to a TAD are combined using Brown’s method, which is meant to combine p-values with possible dependence upon each other^27^. Thus, the combined P-value of genes with non-zero expression in a TAD gene-set is given by

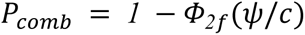

Where 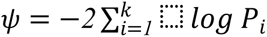 such that *P*_*i*_ is the P-value for the log of non-zero expression of gene i in the single-cell assuming log-normal distribution. The symbol *Φ_2_*_*f*_ represents cumulative distribution function of chi-square distribution with scaled degree *f*. The value of *f* and *c* is calculated using expectation of *ψ* and covariance of P-values to be combined^27^. The combined P-value of genes in a TAD is further adjusted to take into account different artifacts due to bias in gene expression levels and unknown covariates. The P-value adjustment is done with a permutation-based test using transcriptome profile-based combined P-value of TADs of null model^27^. The null model in UniPath was used, which was made by picking the transcriptome of single-cells from multiple studies as well as creating false single-cell expression profiles by taking average values ^27^ of random pairs of cells.

For bulk RNA-seq and microarray-based expression profiles, TADac uses the Gene Set Variation Analysis (GSVA) to calculate gene-set enrichment for genes in TADs for every sample ^28^. GSVA is an unsupervised and non-parametric technique that eliminates the traditional strategy of directly modeling phenotypes within the enrichment score.

For TCGA (https://portal.gdc.cancer.gov/), ^48^ patient data, RNAseq gene expression profiles of 20 cancer types out of 24 cancer types (PANCANCER) were transformed into TAD-activity scores. The data from the Cancer Cell Line Encyclopedia (CCLE)^5^, with 819 samples from different cancer cell lines, were used to calculate TAD-activity scores. In order to estimate change in TAD-activity after developing resistance to drugs, we also used the single-cell expression profiles of HNSCC cancer cell lines published by Sharma *et al.* ^10^ to calculate TAD-activity scores. The gene enrichment scores thus obtained were analyzed for differential activity among resistant and non-resistant cell lines using the Wilcoxon rank-sum test (Supplementary Tables 1-2).

#### Calculating the association between TAD activity and drug-response

A total of 819 CCLE-cancer cell lines were chosen based on the availability of their drug response for 544 drugs in the CTRP^24^ version-2 database. Here, the pIC50 value for drugs was used. The pIC50 value is given as

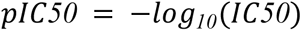

Spearman correlation between TAD-activity scores of cancer lines and their pIC50 value was calculated for every drug.

While modeling the pIC50 value of a drug using machine learning, we used the Lasso regression model. First, we used the ML model with TAD-activity as the feature score. Then, we also used the ML model to estimate pIC50 using only gene-expression values as feature scores. The estimated pIC50 values using the ML model were correlated with actual pIC50 levels and shown in Figure 2D for a few drugs.

#### Calculation of survival P-value

To calculate the significance of the association of TAD-activity and gene expression with survival for TCGA patients, R packages survival and survminer were used ^49^. Input files were TAD-activity scores and clinical data containing days to death, gender, etc, as fetched from the Xena browser ^50^. ‘Status’ column 0 or 1 in clinical data stands for alive and dead, respectively. For calculating the survival P-value for each TAD, the median of each row of GSVA was calculated, and another “median-group” column was created, signifying 1 for GSVA greater than the median and 0 for GSVA less than the median. The clinical information (days-to-death, status (0/1), and patient ID) was merged with GSVA scores and the median-group column. Hence, for each TAD, we created a file from which data was fit in the ‘survfit’ function from R package survival and survminer. Survfit output includes survival P-value for each TAD. In addition, we also calculated the hazard ratio for each TAD using the coxph function.

Different cut-off combinations were implemented to identify TAD significance in survival (high-low), such as 51-49, 60-40, and 70-30 percentiles, to define the group of patients based on the activity of a TAD, in addition to the MaxStat package ^51^. Subsequently, TADs, which were common among such combinations, were selected for further analysis (Supplementary Figure S8, S9). To study the association of patient survival with drug resistance, we created two sets of TADs, i.e., Positive_set and Negative_set, with correlation above 0.3 and below −0.3, respectively. We examined their survival hazard ratio in multiple categories. Furthermore, we also explored the distribution of TADs across different cut-offs of the hazard ratio.

#### Synteny analysis

We obtained the orthologous genes and the corresponding chromosomal coordinates of 69 mammals from Ensembl (https://asia.ensembl.org/index.html). We also downloaded the genome fasta files for the species that were not represented in Ensembl from the DNA Zoo database (https://www.dnazoo.org/) and aligned the genome sequences against that of human genome using last (https://gitlab.com/mcfrith/last). The alignment chain files were then used to map the orthologous positions of genes. We implemented the ‘plotRect’ and ‘plotSegments’ function of ‘plotgardener’ R-package ^52^ to plot the synteny maps from the TSS coordinates of orthologous genes across mammals.

#### Estimation of enrichment of disease/phenotype associated GWAS mutations in EGFR-TAD

We made a list of null model TAD containing random genomic regions of a size similar to EGFR-TAD (chr7:51500000-57100000). The location of GWAS-based mutations was intersected with null-model-based TAD locations and EFGR-TAD to get the count of associated mutations. For a disease X the enrichment of associated mutation in EGFR-TAD was calculated as

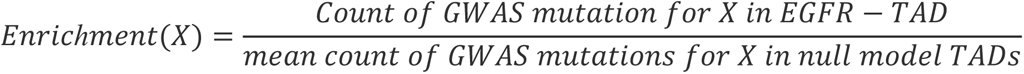

The enrichment scores of top 10 enriched traits and diseases are shown in Figure 6E.

#### Experimental method

The culture of HNSCC patient-derived cell lines (HN137P, HN120P, HN120M, and HN148P) as well as their ChiP-seq and Hi-C based profiling and sequencing, was done at Genome Institute of Singapore.

#### Culture of HNSCC PDC and their cisplatin-resistant model

The creation of the PDC model of HNSCC is described by Chia *et al.* ^26^. Cells were kept in a humidified atmosphere of 5% CO2 at temperature of 37 °C. The cell line identity was authenticated and confirmed by Chia *et al.* through a comparison of STR profile cell lines to their original tumor. The cisplatin version of the same HNSCC cell lines was made by Sharma *et al.* by treating mice xenograft model with cisplatin for 3 weeks. The cells dissociated from the residual xenograft model were isolated into single-cell suspension. Further analysis was performed using Flow-cytometry for cell-type and stem-state markers. FloJo was used for Flow cytometry data analysis.

#### Hi-C profiling

*In situ*, Hi-C was performed according to the method described by Li et al.^53^. Five million cells of HN137, HN120 and HN148 cell lines were cross-linked with 1.5% formaldehyde at room temperature for 10 minutes after which glycine was added to stop the reaction. Lysis of cross-linked cells was performed in 500 µL cold lysis buffer (0.2% (vol/vol) Igepal CA630, 10 mmol/L NaCl, 10 mmol/L Tris-HCl, pH 8.0, mixed with protease inhibitors) on ice for 30 minutes. Washing of pellet performed twice with cold 1× NEBuffer 2, and chromatin was incubated at temperature of 62°C for 10 minutes with in 50 µL 0.5% SDS for solubilization. SDS was quenched using Triton X-100 (Sigma 93443), and digestion of chromatin was done overnight by adding 100 U Mbo I (catalog No. R0147, New England Biolabs) and 25 µL 10X NEBuffer 2. Confirmation of digestion efficiency was done by isolating DNA from a 25-µL aliquot using agarose gel electrophoresis. Filling and labeling of digested ends was done with biotin by adding 37.5 µL 0.4 mmol/L Biotin-14-dATP (catalog No. 19524016, Thermo Fisher Scientific), 1.5 µL of 10 mmol/L dTTP, 1.5 µL 10 mmol/L dCTP, 8 µL of 5 U/µL DNA polymerase I, 1.5 µL of 10 mmol/L dGT and large Klenow fragment (catalog No. M0210, New England Biolabs); and incubated for 90 minutes at temperature of 37°C. A solution of 463 µL water, 120 µL 10X NEB T4 DNA ligase buffer, 12 µL 10 mg/mL BSA, 200 µL 5% Triton X-100 and 5 µL T4 DNA Ligase (catalog No. M0202, New England Biolabs) was added for blunt-end ligation. The solution was incubated for 4 hours at temperature of 16°C. After reversing the cross-link overnight, DNA was purified with the phenol-chloroform method. Next, treatment with T4 DNA Polymerase (New England Biolabs, catalog No. M0203) was performed for biotin removal from unligated ends from Hi-C libraries. To do this, 5 µg Hi-C library was mixed with 2.5 µL of 1 mmol/L dATP, 2.5 µL of 1 mmol/L dGTP and 5 µL of T4 DNA Polymerase (catalog No. M0203, New England Biolabs),10 µL of 10X NEBuffer 2.1, 2.5 µL of 1 mmol/L dGTP, and water was added to a final volume of 100 µL. Incubation of reaction was done for 4 hours at temperature of 20°C and it was stopped by incubating at temperature of 75°C for 20 minutes. Shearing was performed to achieve 400 base pairs (bp) mean fragment size using Covaris M220. Pulling down of biotinylated DNA was done using Dynabeads My One T1 Streptavidin beads (catalog No.65601, Thermo Fisher Scientific). For sequencing library preparation NEBNext Ultra II DNA Library Prep Kit for Illumina was used for. Sequencing was performed using Hiseq 4000.

#### ChIP-seq profiling

Cells were cross-linked for 30 minutes (10% FBS/DMEM containing 1.5% PFA), glycine (1/20 volume of 2.5M) was added and incubation was done on ice (5 min.). It was re-dissolved in a buffer followed by sonication. The buffer used for re-dissolving contained 10 mM Tris-HCl, pH8, 1 mM EDTA, 0.5 mM EGTA, and 1X protease inhibitor cocktail (Roche). For pulling the chromatin, H3K4me3 (04-745, Millipore) or H3K27ac (Abcam ab4729, Lot: 509313) antibody-coupled Dynabeads and 150 μl of ChIP buffer (433 mM NaCl, 4.3% Triton X-100 and 0.43% sodium deoxycholate) were used. For the input control library, 1/10 of the chromatin mixture was kept separately. RIPA buffer was used to wash the beads, which were later incubated at 65 °C for 20 min in 100 μl of elution buffer (50 mM Tris-HCl, pH8, 10 mM EDTA, 1% SDS) with brief vortexing. De-crosslinking of immunoprecipitated chromatin and input samples was done by incubating at temperature of 65 °C overnight and subsequently incubating for 1hour with 0.2 μg/μl of RNase A at temperature of 37 °C followed by treatment with 0.2 μg/μl of proteinase K at 55 °C for 2 hours. DNA was isolated from the chromatin using phenol/chloroform/isoamyl alcohol (49:49:2) and ethanol and precipitated in presence of 40 μg of glycogen. Pellet was then suspended in 10mM Tris-JCL, pH 8 after washing with 70% ethanol. Library preparation and multiplexing for ChIP-seq libraries were performed using the NEBNext ChIP-seq library Prep kit (NEB). ChIP-seq and input control libraries were sequenced using Illumina HiSeq-2500.

#### CRISPR knockout of non-coding regulatory elements in 11q13.3-11q13.4 region

sgRNAs were designed using online CRISPR designing tool “CHOPCHOP” (https://chopchop.rc.fas.harvard.edu/). The sequences for coordinates chr11:71127012-71133012 and chr11:71124803-71130803 were taken from NCBI and given as input sequences for sgRNA designing. Two sgRNA for each coordinate were chosen. The oligos obtained were 20-bp long. Further, complementary pairs for the selected sgRNAs were made using reverse complement tool (https://www.bioinformatics.org/sms/rev_comp.html). The necessary oligos and primers specified by the online tool were ordered from Sigma Aldrich.

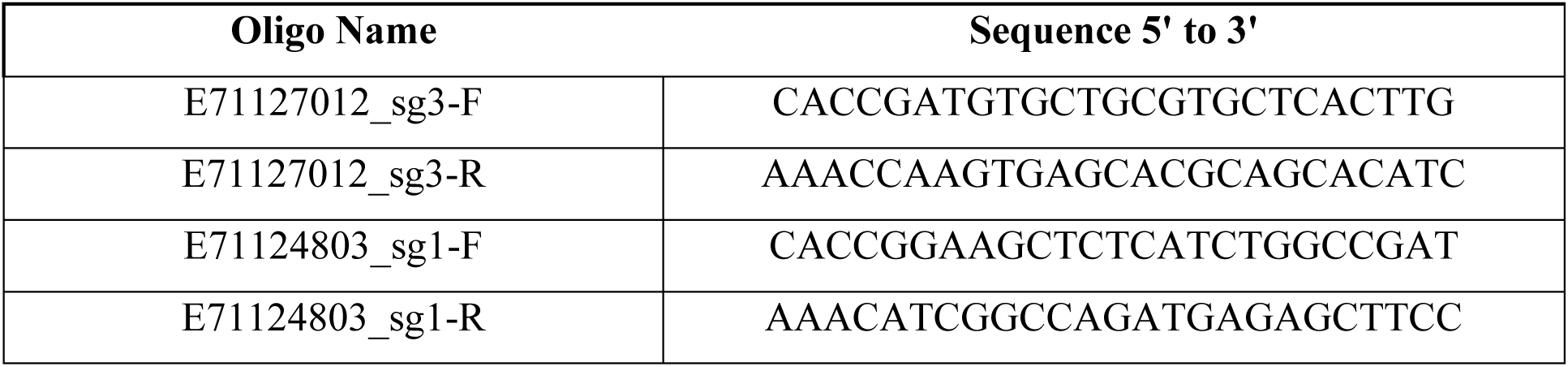

The details about the preparation of sgRNA construct and transfection into cell lines are described in supplementary methods.

## Acknowledgements

We thank A*STAR Singapore as generation of genomic profiles and experiments were funded by Young Investigator grant provided by Biomedical Research council A*STAR ([BMRC/YIG/1510851023] to Vibhor Kumar). We also thank the Bioinformatics Center and Center of Excellence in Healthcare (COEHE) at IIIT Delhi. We also acknowledge the Department of Biotechnology for their support (BT/PR26093/GET/119/11 1/2017). We acknowledge the funding received from Gujarat State Biotechnology Mission (GSBTM) (GSBTM/JD(R&D)/618/21–22/1222 to Amit Mandoli) and Indian Council of Medical Research (ICMR), India (5/13/50/2022/NCD-III to Amit Mandoli).

## Author’s contribution

V.K and R.D. conceived the project. N.P., M.S. performed analysis for TAD based analysis. While I.P.J., O.C. and S.M., A.M.1 contributed to basic analysis and figure generation. A.S. and M.H. were involved in cell culture of HNSCC cells. C.G.A. and M.H. performed Hi-C and ChIP-seq experiments with support and guidance from R.F, R.D and A.S. CRISPR knockout experiments were performed by P.P. and A.M.2.. J.B. and K.S. performed analysis for tracking synteny of genes in TADs. N.P., M.S. and V.K. were involved in writing of manuscript while V.K. also supervised the project.

## Data availability

The Hi-C and ChIP-seq profiles generated by this study can be downloaded from GEO database (GSE249524, https://www.ncbi.nlm.nih.gov/geo/query/acc.cgi?acc=GSE249524 Reviewer’s access code: **cdwnaiqefzktjih**).

Hi-C, ChIP-seq, and single-cell profiles published by other studies were also used here. ChIP-seq profiles of HN120M cell lines were adopted from data-set with GEO ID: GSE120634. While the Hi-C profile of Cardiomyocytes was downloaded from the GEO dataset (GSE106690, 4DNFIN39NO4O.hic from 4dnucleome repository, https://data.4dnucleome.org/files-processed/4DNFIN39NO4O/). The Hi-C profile of Astrocytes was downloaded from 4dnucleome database (https://data.4dnucleome.org/files-processed/4DNFITPO1WTY/). Additional TAD boundaries were downloaded from the database TADKB (http://dna.cs.miami.edu/TADKB/).

The single-cell expression profiles of HNSCC cell lines HN120M, HN120MCR, HN137P, HN137PCR, HN137M, and HN137MCR are present in the GEO database (GSE117872). The bulk expression profile of cancer cell lines in the CCLE database (https://sites.broadinstitute.org/ccle/datasets). The IC50 values for cancer cell lines are available at the CTRP website (https://portals.broadinstitute.org/ctrp.v2.1/).

All the result data-set (including correlation values, association P-value, etc) obtained from our codes have been uploaded to the link http://reggen.iiitd.edu.in:1207/tads/TADCancer/datasets/ (username: reggen, password:unipath@123)

## Code availability

The codes for TADac and figures in this study can be downloaded from the GitHub link: https://github.com/reggenlab/TADs_repository. DFilter can be downloaded from https://reggenlab.github.io/DFilter/.

